# Enzymatic evolution driven by entropy production

**DOI:** 10.1101/319814

**Authors:** A. Arango-Restrepo, J.M. Rubi, D. Barragán

**Affiliations:** Institution, Address

## Abstract

We show that the structural evolution of enzymes is largely influenced by the entropy produced in the enzymatic process. We have computed this quantity for the case in which the process has unstable and metastable intermediate states. By assuming that the kinetics takes place along a potential barrier, we have found that the behavior of the total entropy produced is a non-monotonic function of the intermediate state energy. By diminishing the number of metastable intermediate states, the total entropy produced decreases and consequently the enzyme kinetics and the thermodynamic efficiency are enhanced. Minimizing locally the total entropy produced for an enzymatic process with metastable intermediate states, the kinetics and the thermodynamic efficiency are raised. In contrast, in the absence of metastable intermediate states, a maximum of the entropy produced results in an improvement of the kinetic performance although the thermodynamic efficiency diminishes. Our results show that the enzymatic evolution proceeds not only to enhance the kinetics but also to optimize the total entropy produced.

## INTRODUCTION

Enzymes are the devices responsible for all the energy conversions in the cell. They are very elaborately adapted for this function and as catalysts they have the capacity to accelerate the rate of biochemical reactions (1). It has been shown that enzyme performance depends on their structure, reactivity, catalytic efficiency and activation energy. However the thermodynamic efficiency of the enzyme activity is related to the kinetic performance is currently an open problem.

SThe description of enzymatic mechanisms demands the use of a very intricate standard free-energy landscape containing multiple minima and transition states (2). However, there is experimental evidence (3) showing that enzymatic processes take place along just one path (see Fig. 1). For instance, processes mediated by adenylate kinase (AdK) arise along only one possible path having multiple intermediate conformations that may facilitate a rapid transition (4). It has been argued that relatively small free-energy differences between conformations could improve the fine control of transitions by environmental perturbations and signaling (5). Additionally, a recent study of the enzyme mechanisms has shown that on average each reaction undergoes 4.3 steps and involves 2.7 intermediates (6).

**Figure 1:**
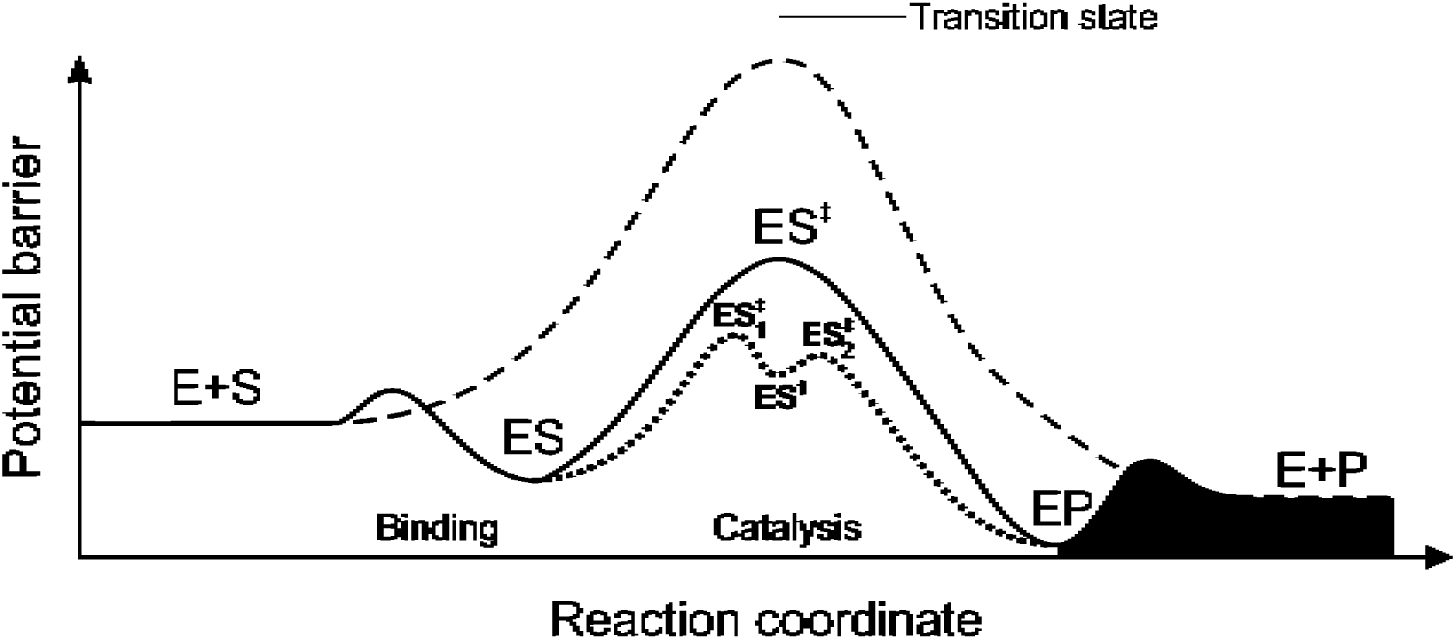
Potential barrier for a general enzymatic process: Free energy landscape for a complete enzymatic process; substrate binding *E* + *S* ⇌ *ES*, catalysis *ES* ⇌ *EP*, product release *EP* ⇌ *E* + *P*. Dashed line corresponds to a non-enzymatic process *S* ⇌ *P*. The catalysis takes place in the catalytic step in which we can find the transition states. The intermediate states that can be unstable (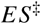, 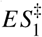 and 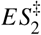) or metastable (*ES*^†^).

The catalytic nature of the enzyme has been related to an electrostatic stabilization of the transition state during the enzymatic process, as shown in the catalysis zone in Fig. 1. This electrostatic effect can be seen as a combination of the reduction in both the energy of the transition state and in the reorganization energy (7). The active sites are pre-organized in a geometry which minimizes conformational reorganization and maintains an optimal stabilization for the transition state in each step during catalysis (8, 9). For instance, theoretical results for serine esterases show that these perform multiple steps minimizing the conformational rearrangements to avoid energetic conflicts following a “minimally frustrated” path or a minimum energy path.(7, 10).

How proteins and enzymes evolve remains as an open question (11). Average enzymes exhibit moderate catalytic efficiencies (12) because maximal rates may not evolve in cases that can be associated with weaker selection pressures (13). Also, it seems likely that the catalytic efficiency of some enzymes toward their natural substrates could be increased in many cases by natural or laboratory evolution (12). Evolutionary pressures play a key role in shaping enzyme parameters (related to potential barrier); that is, maximal rates have not evolved in cases where a particular enzymatic rate is not expected to be under stringent selection (12). Initially, it is proposed that enzymes have evolved firstly to optimize the speed of the actual breaking-bonding chemical processes, optimizing the potential barrier for the diffusion through transition states to finally tune the rates (7, 14). However, evolutionary pressures have their limits and cannot enhance the kinetics of an enzyme at its optimum or ad infinitum (12).

Enzyme evolution seems to be globally driven by physiological conditions to improve reaction rates by means of changes in the structure to stabilize the transition states thus decreasing activation energies and modifying the number of metastable intermediates. However, most of the enzymes are far from optimal kinetic conditions not only due to evolutionary aspects and physical-chemical constraints because of thermodynamic restrictions. There is no clear evidence of the fact that improving reaction rates leads to an improvement of thermodynamic efficiency of catalytic processes. As far as we know, enzymes must be stable in order to conserve their structure and locally unstable in order to function. Therefore, there is a debate about which trajectory an enzyme should take to evolve structurally and why these trajectories are taken. In that sense, thermodynamically, the evolution of the enzyme could be conditioned to decrease or increase the entropy produced in the catalytic step of the process.

Our purpose in this article is to analyze the transition between enzyme-subtract and enzyme-product states (the catalytic step of the enzymatic process of Fig. 1). As the enzyme activity takes place at the mesoscopic level (15), the enzymatic processes is modeled by a diffusion through a potential barrier whose intermediate states are parametrized by means of a reaction coordinate, in which the shape of the potential barrier is related to the structural changes of the enzyme (16). Therefore, we use the Mesoscopic Non-Equilibrium Thermodynamics (MNET) (8, 17) formalism to quantify the entropy produced in the transit/diffusion through the potential barrier and thus obtain the reaction rates defined in the space of the reaction coordinate (17). By modelling the potential barrier with and without local minima, we compute the entropy production. Here, we present how the enzyme evolution could be directed towards an optimization of the total entropy produced as a function of the activation energy restricted to the relaxation time.

## METHODS

MNET provides a general framework for the study of small-scale far from equilibrium systems which unify thermodynamic and kinetics aspects (8, 17, 18). Activated processes such as chemical reactions, adsorption (19), electro-chemistry (20), unzipping RNA (21), active transport (15) and biochemical cycles (22) that take place in physico-chemical and biological systems, can be viewed as processes that proceed along an internal coordinate that parameterizes the different molecular configurations. Considering that the catalytic efficiency of enzymes depends on their structures (23), we will adopt a molecular perspective to analyze the process through the MNET formalism. Moreover, we will consider that the rearrangement of the enzyme active site in a process determines the activation energy and the reaction rate (24).

### The model

In the process, an enzyme-substrate complex transforms into an enzyme-product complex in a homogeneous close system under isothermal conditions. The transformation takes place along the reaction coordinate *γ* defined from 0 to 1. The evolution of the probability density to find the complex in the state *γ* at time *t* is governed by the continuity equation:

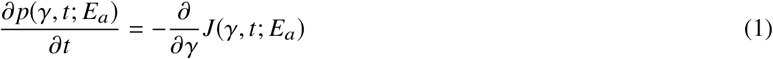

Here, *J* is the current along the reaction coordinate and *E_a_* the activation energy. Assuming local equilibrium along the *γ*-coordinate, we use the Gibbs relation and the continuity equation (Eq.(1)) to obtain the entropy production of the enzymatic process at the mesoscale (17):

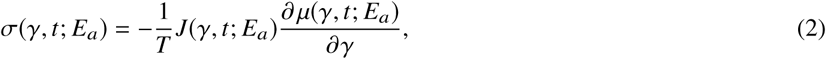

where *µ* the chemical potential. From the entropy production (Eq.(2)), we can write the linear law for the current, given by

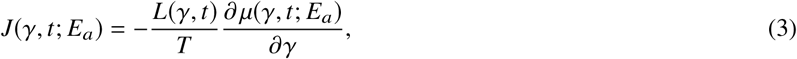

whith *L* an Onsager coefficient which is in general a function of the state of the enzyme and the time. Using the fact that the Onsager coefficient is in very good approximation proportional to *p*(*γ*, *t*; *E_a_*), we introduce the constant diffusion coefficient 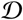 through the expression:

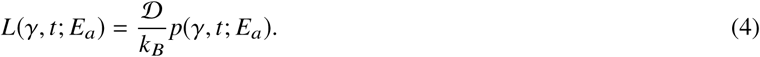

The chemical potential along the *γ*-coordinate is given by

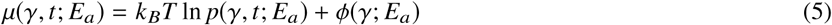

where *ϕ* is the potential barrier of enthalpic nature through which the diffusion process takes place along the *γ*-coordinate. This barrier is a function of the activation energy and is intimately related to the enzyme structure. By inserting the expression of the current (Eq.(3)) into the continuity equation (Eq.(1)), we obtain the Fokker-Plank type kinetic equation:

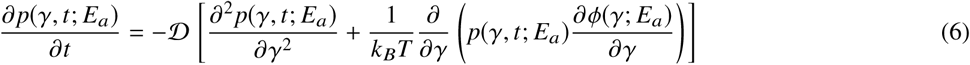

By solving this equation, we can compute the entropy production rate and from this, the total entropy produced in the process

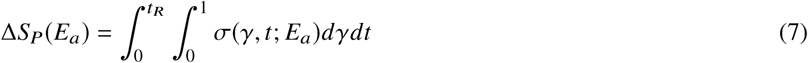

where *t_R_* is the relaxation time of the process. To study the thermodynamic efficiency, in the following sections we will analyze the lost work defined as *W_L_* = *T*Δ*S_P_*(*E_a_*).

The diffusivity through the potential barrier 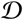 is the only kinetic parameter of the model. This parameter depends neither on the activation energy nor on the shape of the potential barrier and it will be used to construct a dimensionless time: 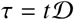. As we will show in the results section, for the fastest processes (with activation energy ∝ 1*k_B_T*) the dimensionless time is around 1. Considering the time reported in (25), the diffusivity could take a reference value of 10^14^*s*^−1^.

### Modeling potential barriers

Enzymes can be modified artificially by changing the potential barrier landscape of the enzymatic process. For instance, although triose phosphate isomerase is considered an advanced and perfect enzyme, its structure was modified by changing the activation energy of the enzymatic process thus altering the reaction rate. These modifications of the energy landscape having an impact on the kinetics of the process could be the next step in the kinetic evolution of the enzyme (26). In this section, we present particular forms of the energetic barriers and in the results section we analyze the lost work as a function of the parameters shaping the barrier.

Two particular forms of the barriers are shown in Fig. 2(a) and their modifications are depicted in Fig. 2(b)-(d). Barriers with an unstable intermediate are shown in Fig. 2(b) for different activation energies. Barriers with a metastable intermediate are shown in Figs 2(c)-(d) for different intermediate energies and transition-state energies. These barriers will be used in our numerical calculations to study enzymatic reactions taking place in a closed system. Notice that in Figs. 2(c)-(d) the difference between the maxima 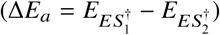 which is related to the activation energy is fixed. The proposed potential barriers are bistable and tristable functions modelled by polynomials (see the supplementary information). They must fulfill some thermodynamics conditions. For instance, at *γ* = 0 and *γ* = 1 the values of the potential coincide with the standard chemical potential of the *ES* and *EP* states. Moreover, barriers representing a process with only unstable states there is only one maximum corresponding to the transition state. For those with a metastable intermediate state there are two maxima and one minimum representing the transition state and the metastable state. Finally, the macroscopic equilibrium relations must be fulfilled (see supplementary information).

**Figure 2:**
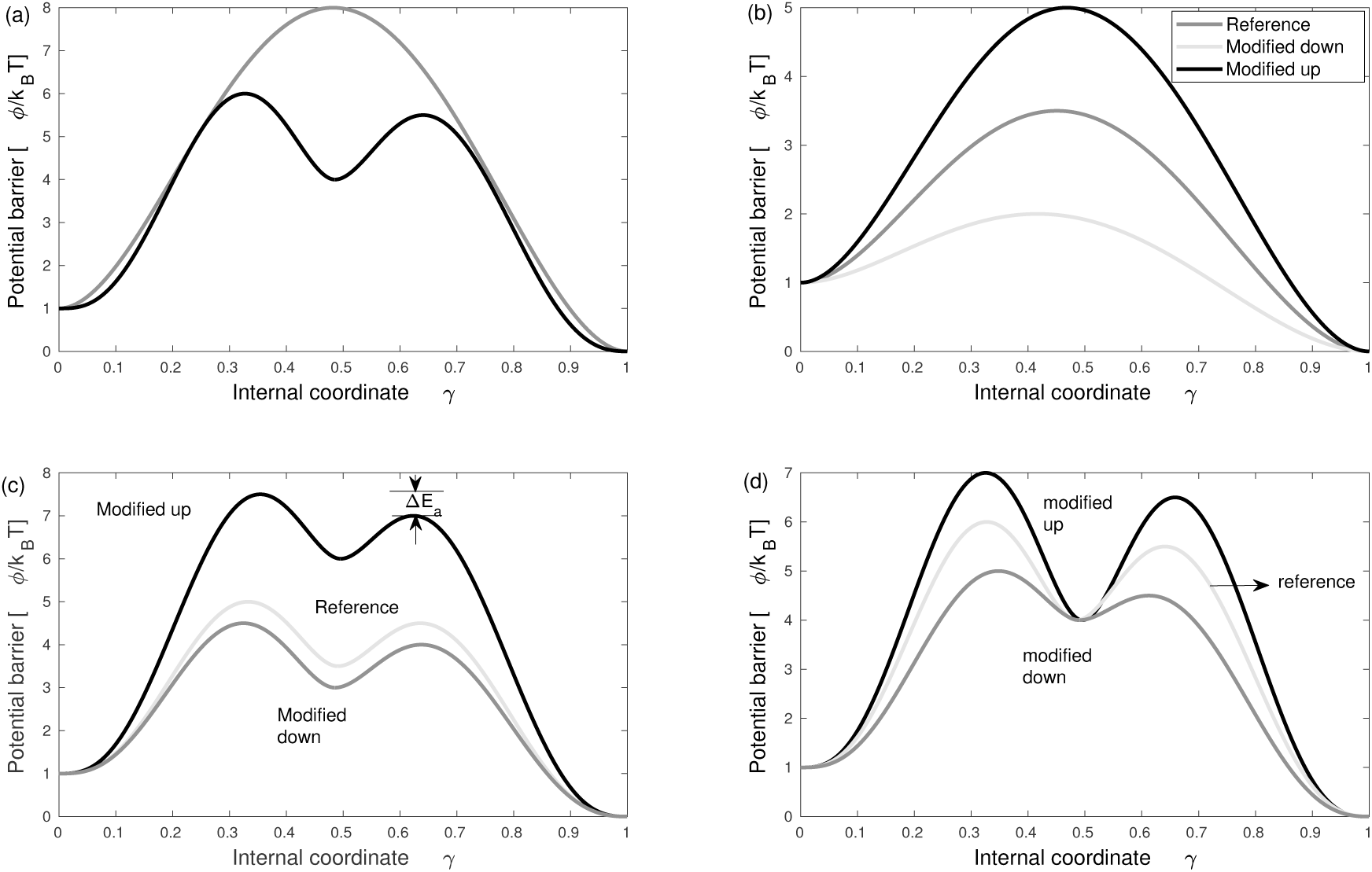
Potential barrier *ϕ*(*γ*) as a function of the reaction coordinate *γ* for metastable and unstable intermediates. Potential barrier for: (a) an unstable intermediate state (gray line) and a metastable intermediate state (black line) in an enzymatic process 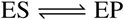. (b) an enzymatic process with unstable intermediate state for three activation energies due to modifications or mutations. (c) a metastable intermediate state for three intermediate energies due to modifications or mutations, for Δ*E_a_* fixed. (d) a metastable intermediate state for fixed intermediate energy but different activation energies due to modifications or mutations, for Δ*E_a_* fixed.

Enzymes might mutate to enhance their functions. We have recognized two kind of barriers in which enzymatic processes takes place. The way potential barriers are modified, shown in Fig. 2, may mimic the enzyme mutation process. Fig. 3 exemplifies a scenario where the enzyme through mutations could change the energy of the intermediate states in the catalytic process. The proposed mutation factor *ζ* depends on the activation energy and on the changes of the potential barrier with the energy of the intermediate states. An energy landscape (in terms of the barriers) as a function of the mutation factor (characterizing possible mutation paths) and the reaction coordinate is given by,

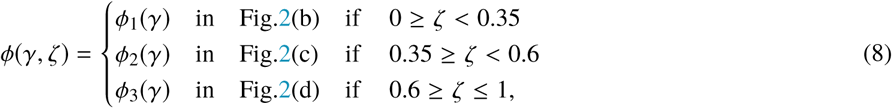

The mutation factor helps us exemplify a hypothetical landscape of barriers in an enzymatic process. Notice that there are more possible ways to change the energy of the intermediate and transition states and therefore the shape of the potential barrier. Nevertheless, here we illustrate some representative cases as a first approximation.

**Figure 3:**
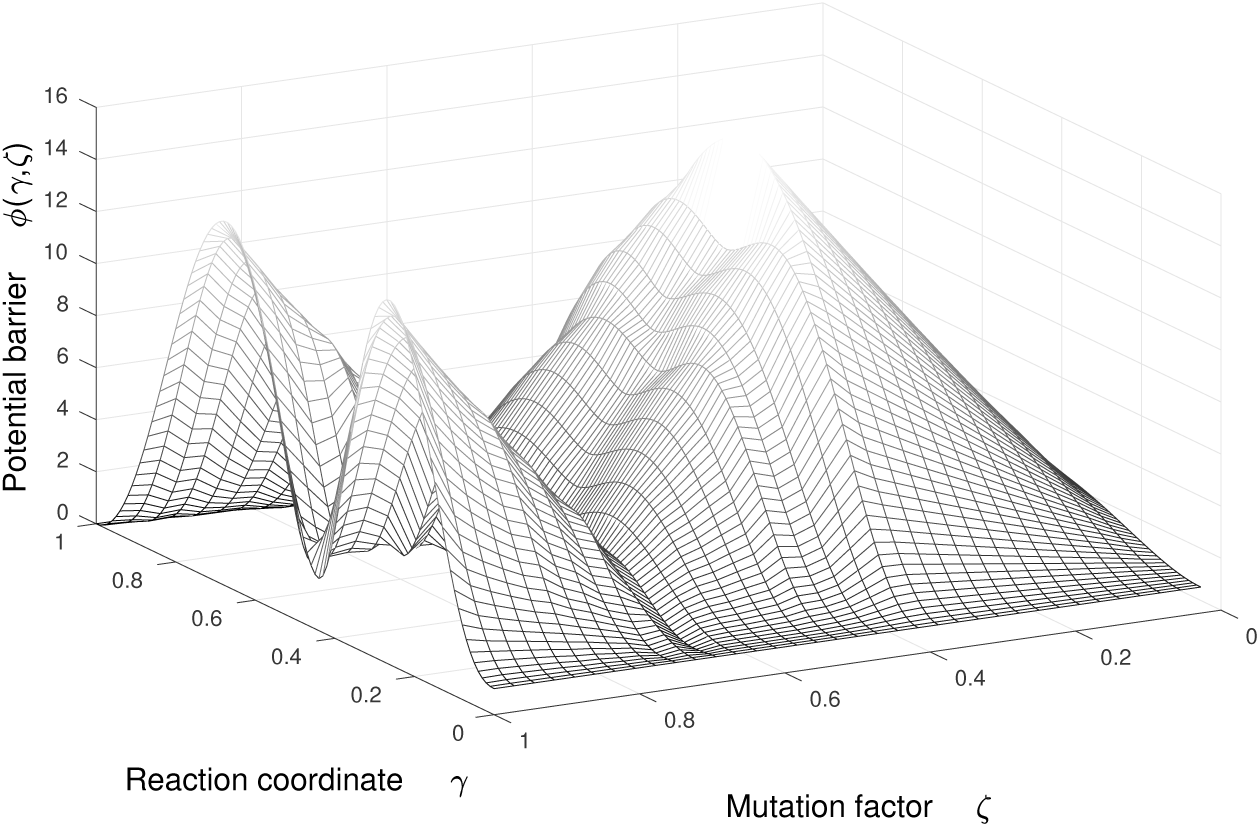
Potential barrier *ϕ*(*γ*) as a function of the reaction coordinate *γ* and of the mutation factor *ζ*. Sketch of a generic potential barrier representing different possible energetic configurations of the enzymes related to their structural configuration and parametrized by *ζ*.

Why nature uses mutations to increase or decrease the energy of intermediate states (or the activation energies) is as of yet unanswered. We hypothesize that not only kinetics but also energetic issues are involved in enzymatic mutations. Under this perspective, the lost work related to the more energetic states in an enzymatic process (transition states), is key to understanding how evolution could enhance enzyme kinetics.

## RESULTS AND DISCUSSION

The entropy production rate calculated by using the potential barrier shown by the gray line in Fig. 2(a) is plotted as a function of dimensionless time in Fig. 4. The curves correspond to different values of the activation energy. The relaxation times *t_Ri_* are indicated by arrows, and are proportional to the activation potential energy (*ϕ*^*^− *ϕ*(0)/*k_B_T*). A decrease of the activation potential energy, however, leads to an increase of the enzymatic reaction rates and to an increment of the initial entropy production. In this sense, an eventual kinetic principle that predicts infinite rates will also predict an infinite entropy production, which seems to be meaningless. Since the dynamics of the enzyme can be characterized by the entropy generation and the relaxation time, the total entropy produced is in general a non-trivial function of the activation energy and can give us information about which are the most efficient configurations of the enzyme.

**Figure 4:**
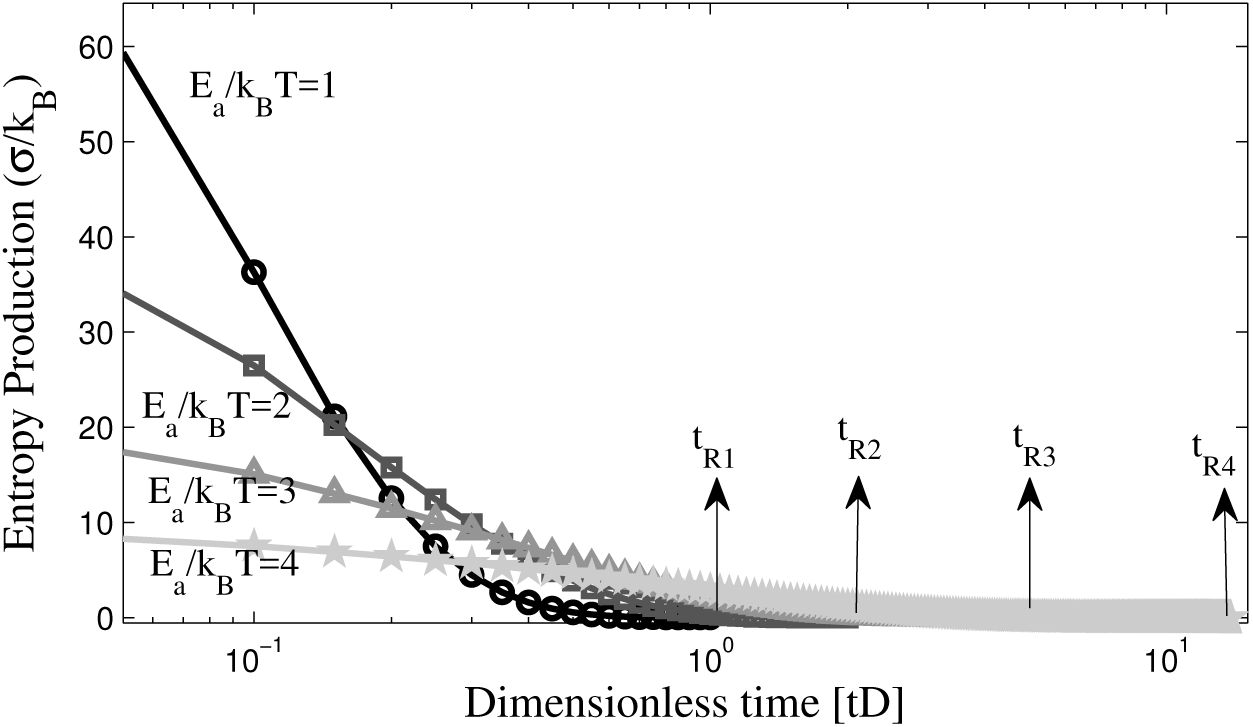
Entropy production *σ*(*E_a_*) as a function of the dimensionless time 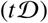 for different activation energies, for processes with unstable intermediates. The dimensionless time axis is presented in logarithmic scale.

Integrating the entropy production over *γ* and over time for the cases shown in Fig. 4, we find that lost work always decreases for increments of the activation potential energy and of the relaxation time of the process (Fig. 5, black dashed line). One could then infer that for a given process a maximization of the lost work (or a minimization of the thermodynamic efficiency) results in an enhancement of the kinetics and a minimization of the relaxation time. This case will be considered as the reference scenario. Nevertheless, a more complex behavior of enzymatic processes with metastable intermediate states can be found in two ways.

**Figure 5:**
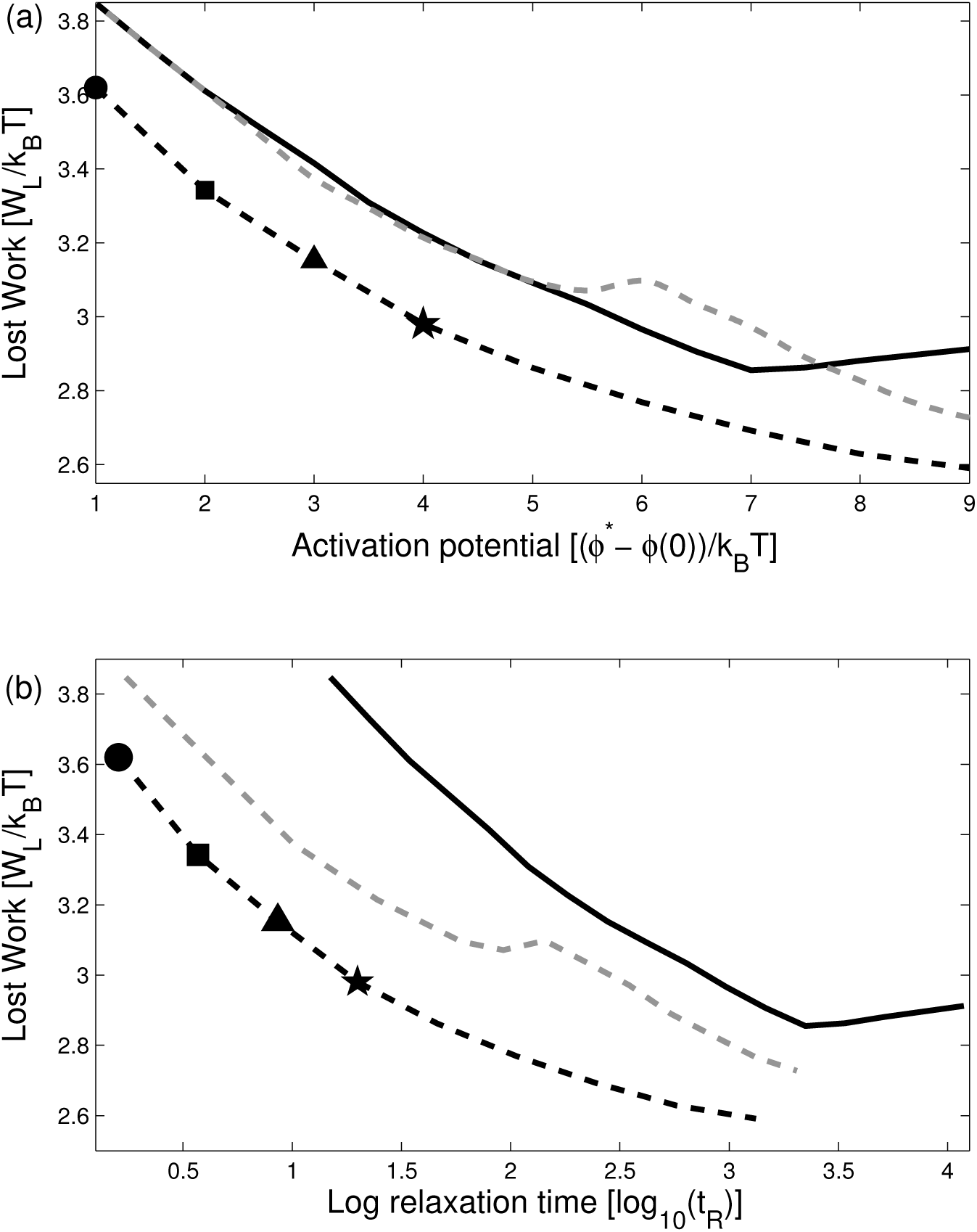
(a) Lost work as a function of the activation potential energy for enzymatic process. (b) Lost work as a function of the relaxation time of the process. Integration over time for each data (*E_a_*/*k_B_T* = 1; 2; 3; 4) shown in Fig. 4 gives us lost work as a function of the activation energy (or relaxation time) for a process only with unstable intermediate states (black dashed line), as shown in Fig. 2(b). Lost work for a process with metastable intermediates states: intermediate state energy is fixed (black continuous line) forming a deep well in the potential barrier for Δ*E_a_* fixed, as shown in Fig. 2(d); intermediate state energy increases proportional to the first activation energy (gray dashed line), for Δ*E_a_* fixed, as shown in Fig. 2(c).

First, by changing the shape of the potential barrier, as shown in Fig. 2(c), and by computing the lost work for different values of the activation potential energy of the metastable state (*ES*), in which Δ*E_a_* is fixed, we obtained the lost work shown by the gray dashed line in Fig. 5. Lost work exhibits a local minimum (in this case around 5*k_B_T*) and when it approaches this value, it starts to decrease until it converges with the value found in the reference scenario (a process with only unstable intermediates). This could be plausible because for high intermediate energy, the condition to overcome the barrier is similar to that of the case shown in Fig. 2(b) for high intermediate energies. Moreover, the relaxation times are similar to those of the reference scenario because the energies of transition states and the metastable intermediate are alike. Therefore, the process cannot be divided into two subprocesses, as in the case shown in Fig. 2(d), giving rise to a slowing down of the kinetics.

In a second way, by changing the shape of the potential barrier, as shown in Fig. 2(d) for different energy values of the transition states 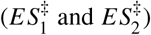, we obtained the lost work shown by the black continuous line in Fig. 5. In Fig. 5(a), there is a local minimum (in this case around 7*k_B_T*) and from this value the lost work increases. Notice that for higher values of the activation potential, the height of the two peaks (transition states energies) and the depth of the well in the potential barrier increases. In this case, the probability density around the metastable state at the beginning of the process is low and consequently goes up considerably, increasing the rate of jumps over the second wall of the potential barrier. This increases the relaxation time and gives rise to the appearance of two stages in the enzymatic process, contrary to what happens at low values of the transition states as shown in Fig. 2(c). Therefore, depending on the value of the activation potential, we can observe one-stage or two-stages processes. The minimum observed in Fig. 5 reveals the presence of a transition between both types of processes.

From Fig. 5 we conclude that in the most efficient processes taking place in a system with a metastable intermediate state, the energy is similar to that of the transition states to avoid deep wells in the potential barrier. One can thus expect that if the enzymatic process involves a metastable intermediate state (as in Fig. 2(c)(d)), the enzyme could evolve towards structures such that the intermediate state involved have an energy similar to that of the transition states in order to decrease the presence of subprocesses that may impair the thermodynamic and kinetic efficiencies.

Additionally from Fig. 5 we conclude that a given process with metastable intermediate does not necessarily maximize the lost work to enhance the kinetics. It is found that in the surrounding of the local minimum, kinetics and energetics of the process could be simultaneously improved. Therefore, by decreasing lost work we obtain an enhancement of the kinetics. One can thus expect that if the enzymatic process involves a metastable intermediate, the enzyme could evolve towards a structural configuration such that the process has an activation potential energy that ensures a local minimum in lost work.

Notice that the lost work for an enzymatic process with a metastable intermediate (Fig. 2(c)-(d)) is larger than for a process with unstable intermediates (Fig. 2(b)). This is because of the presence of two (or more) sub-processes involved to overcome the potential barrier. Therefore, an additional scenario could exist in which the enzyme may evolve towards a more efficient structural configuration. Thus enzymes catalyzing processes with metastable intermediates could increase the energy of these intermediates and shift the potential barrier to the ones shown in Fig. 2(b), thereby avoiding sub-processes. In this scenario, once the enzymatic processes avoids the metastable intermediate state, the enzyme may evolve towards a configuration in which the activation energy decreases in order to maximize the lost work, as indicated in Fig. 5.

## CONCLUSIONS

In this article, we have shown how enzyme evolution is conditioned to an optimization of the total entropy produced as a function of the activation energy. We have found that the shape of the potential barrier determines the thermodynamic efficiency of the activated processes because lost work depends on the nature of the intermediate state.

An increase of the reaction rate and of the thermodynamic efficiency can be induced by decreasing the number of intermediates states or by decreasing the energetic difference between intermediate and transition states. This conclusion is in accordance with the observation that some enzymes minimized the number and height of the peaks of the energetic barriers between consecutive conformations (10). Moreover, our conclusion is in accordance with a minimum energetic path concept (“minimally frustrated” path)(7).

In the framework of the transition state theory, the efficiency of an enzyme has been related to the number of conformational changes that it must undergo to return to its original state. From this perspective, if the energy related to the conformation changes is minimized, higher reaction rates could be obtained (27). In the context of the present work, however, if the energy related to conformational changes is minimized, lost work is minimized as well since it entails a decrease of the energetic barriers, thus avoiding metastable intermediates. Therefore, the energy related to conformational changes is not minimized to improve the kinetics (as claimed in the literature (27)) but rather to optimize the thermodynamic performance of enzymatic process.

Our results support the idea that enzymatic process have on average 2.7 intermediates per reaction (6) and not just one or even no intermediate at all. This is because the structural configuration of the enzyme is determined by a local minimum in lost work of the enzymatic process as a function of the activation potential energy. In this scenario, an enhancement of the kinetics around the minimum leads to a decrease of the lost work and consequently to an improvement of the thermodynamic efficiency, thus favoring the enzyme’s structural stability and the catalytic activity.

The fact that central metabolism enzymes are more efficient than secondary metabolism enzymes could be explained from a thermodynamic point of view. This is because the evolutionary pressures (12, 28) may be related to a high probability of modifications in the enzyme structure which could changes the activation energies. This forces these enzymes to evolve either towards a local minimum of lost work thus enhancing their kinetic and thermodynamic efficiency, or towards a maximum of lost work to improve only their kinetics.

The model presented here might be used to analyze the effect of temperature and pH (considered as the mutation factors) over kinetics through the modification of the shape of the potential barrier upon variations of those quantities.

## AUTHOR CONTRIBUTIONS

All authors have contributed equally developing the research idea, data analysis, and manuscript redaction. Author 1 carried out all the simulations.

## ACKNOWLEDGMENTS

The authors thank the Facultad de Ciencias of Universidad Nacional de Colombia Sede Medellin for partial supporting under the grant QUIPU 201010021328. Further, this work has been supported by MINECO of the Spanish Government under Grant No. FIS2015-67837-P.

## SUPPLEMENTARY MATERIAL

Supplementary information to this article can be found in the attached pdf.

